# The MHC class-II HLA-DR receptor mediates bat influenza A-like H17N10 virus entry into mammalian cells

**DOI:** 10.1101/507467

**Authors:** Efstathios S Giotis, George Carnell, Erik F. Young, Saleena Ghanny, Patricia Soteropoulos, Wendy S Barclay, Michael A Skinner, Nigel Temperton

## Abstract

Bats are notorious reservoirs of diverse, potentially zoonotic viruses, exemplified by the evolutionarily distinct, influenza A-like viruses H17N10 and H18N11 (BatIVs). The surface glycoproteins [haemagglutinin (H) and neuraminidase (N)] of BatIVs neither bind nor cleave sialic acid receptors, which suggests that these viruses employ cell attachment and entry mechanisms that differ from those of classical influenza A viruses (IAVs). Identifying the cellular factors that mediate entry and determine susceptibility to infection will help assess the host range of BatIVs. Here, we investigated a range of cell lines from different species for their susceptibility to infection by pseudotyped viruses (PV) bearing bat H17 and/or N10 envelope glycoproteins. We show that a number of human haematopoietic cancer cell lines and the canine kidney MDCK II (but not MDCK I) cells are susceptible to H17-pseudotypes (H17-PV). We observed with microarrays and qRT-PCR that the dog leukocyte antigen DLA-DRA mRNA is over expressed in late passaged parental MDCK and commercial MDCK II cells, compared to early passaged parental MDCK and MDCK I cells, respectively. The human orthologue HLA-DRA encodes the alpha subunit of the MHC class II HLA-DR antigen-binding heterodimer. Small interfering RNA- or neutralizing antibody-targeting HLA-DRA, drastically reduced the susceptibility of Raji B cells to H17-PV. Conversely, over expression of HLA-DRA and its paralogue HLA-DRB1 on the surface of the unsusceptible HEK293T/17 cells conferred susceptibility to H17-PV. The identification of HLA-DR as an H17N10 entry mediator will contribute to a better understanding of the tropism of the virus and will elucidate its zoonotic transmission.

## Main

Outbreaks of SARS, MERS, Nipah and Ebola have highlighted the critical need to focus on the zoonotic potential of known, and novel, bat viruses to improve forecasting, prevention and control of epidemics. Viral diversity in bats is exemplified by the discovery of the enigmatic influenza A-like viruses (BatIVs) H17N10 and H18N11 in asymptomatic New World bats (*Sturnira lilium* and *Artibeus planirostris* respectively)^1^, ^2^ and more recently by the detection of a virus related to avian H9N2 in Egyptian *Rousettus aegyptiacus* bats^3^. Such discoveries prompted investigation of the pandemic potential of these viruses and led to concern that bats may be a neglected reservoir of novel influenza viruses^4^.

Influenza A viruses (IAVs) are enveloped orthomyxoviruses with eight single-stranded negative-sense viral RNAs (vRNAs) encapsidated into viral ribonucleoproteins (vRNPs). The original source of classical IAVs is aquatic birds, from which they emerge, *via* genome reassortment and mutation, to cause sporadic pandemics in humans, lower animals and other birds^5^, ^6^. They are classified into different subtypes based on their envelope glycoproteins (trimeric haemagglutinin, HA: H1-H18, and tetrameric neuraminidase, NA: N1-N11)^7^. HA is synthesised as a precursor protein in infected cells and its cleavage by host cell proteases sets in motion a complex series of events that is initiated by receptor binding and is terminated with the penetration of the virus into the cytoplasm of target cells^8^. HA conventionally attaches to host-specific sialic acid (SA) moieties^6^. These are terminal sugars of larger carbohydrate chains attached to the cell membrane by the lipids or proteins that they decorate. When HA attaches to them, it triggers endocytosis of the virus into membrane bound endosomes^9^, ^10^. Acidification of the endosome induces conformational changes to HA, which lead sequentially to: insertion of the hydrophobic “HA fusion peptide” into the host membrane, juxtaposition of the viral and endosomal membranes and subsequent release of the vRNPs into the cytoplasm via a fusion pore^11^, ^12^. In contrast, NAs are glycosidases which primarily cleave cell surface SA and therefore facilitate the release and spread of virus progeny upon egress, as well as disaggregation of virions before entry^13^, ^14^.

The crystal structures of BatIV HAs and NAs revealed divergence of their protein conformations from those of conventional IAVs, suggesting distinct binding and functional properties^2^, ^15^, ^16^, ^17^. Bat H17 and H18 proteins have typical HA folds but lack an obvious cavity to accommodate SA^2^, ^15^, ^17^. The cell receptors for the BatIVs are as yet unidentified, but they are clearly not SA moieties, a conclusion reached by several studies^15^, ^18^, ^19^. Furthermore, bat N10 and N11 are structurally similar to classical NAs but lack conserved amino acids for SA binding and cleavage^2^, ^16^, ^17^ and do not exhibit typical neuraminidase activity^18^, ^20^. Initial efforts to isolate infectious BatIVs directly from bats have failed, mainly because the receptors were unknown and susceptible cell lines were unavailable^21^, ^22^, ^23^. Attempts to circumvent these limitations have included H17- or H18-pseudotyped vesicular stomatitis virus (VSV^19^, ^24^), engineered BatIV/IAV chimeric viruses^23^, ^25^, and authentic BatIVs reconstructed using reverse genetics^26^. H17-VSV was able to infect bat cell lines (EidNi, HypNi, and EpoNi) but only a few, common cell lines of flightless mammals, including some of human (U-87 MG glioblastoma and SK-Mel-28 melanoma) and canine (RIE1495 and MDCK II kidney) origin^19^, ^24, 26^. These cells could also be infected with reconstructed H17N10 and H18N11 viruses^26^. The ability of BatIVs to infect mammalian cell lines *in vitro*, and their unconventional features, raised concerns about their zoonotic and epidemic potential. Identifying the BatIV cell surface receptors and delineating the mechanistic basis of the host-virus interaction are key to assessing their potential host range and public health significance.

HIV-1 derived-pseudotypes bearing heterologous envelope proteins (PV) have been used widely for the assessment of cellular tropism and the identification of cellular receptors or attachment factors for a range of viruses^27^, ^28^, ^29^, ^30^, ^31^. Such pseudotypes have proved a reliable model to study the capacity of H17N10 for entry into various cell lines^27^, ^32^. Using this approach, we have shown that H17-PV, and H17N10-PV (A/little yellow-shouldered bat/Guatemala/060/2011) are recovered from producer HEK293T/17 cells exclusively in the presence of the human airway trypsin-like protease (HAT) or the transmembrane protease, serine 2 (TMPRSS2) (Fig. 1a)^32^. In this study, a panel (*n*=35; Supplementary material 1) of cell lines from different tissues and species were challenged with H17- and/or N10-PV to study the distribution of the H17N10 receptor(s) (Fig. 1b). Efficiency of infections with PV was quantified (after 48 h) by the expression of a firefly luciferase (FLuc) reporter gene encoded by the lentiviral genome. Parallel infections were conducted with PV bearing either classical H5 (H5-PV; A/Vietnam/1194/2004; H5N1 clade 1) or VSV-G (VSV-G-PV) glycoproteins (the latter displaying very broad tropism) as positive controls to eliminate possible post-binding blockage factors. HIV particles produced in the absence of a viral envelope protein (Δ-env) served as a negative control. H17-PV displayed highly limited host and species cell tropism, suggesting that the H17 cellular receptor(s) are not ubiquitous (Fig. 1b). Of note, the bat cell lines (lung & kidney) from *S. lilium* in which H17N10 was discovered, were not susceptible to PV. This implies that expression of the H17-putative receptor(s) and/or viral entry-related host factors was either lost during immortalisation or is tissue-type restricted. Conversely, we show that the dog epithelial kidney MDCK II (unlike MDCK I) cells are susceptible to H17-PV with titers comparable to those of control VSV-G- and H5-PV in the range of 10^6^ to 10^7^ RLU/ml (Fig. 1c). They were not susceptible to PV expressing N10 alone and co-expression of N10 with H17 did not improve infection of MDCK II (Fig. 1c), which suggests that N10 has a dispensable role in viral entry (*in vitro*). To characterise the H17 putative receptors, MDCK II cells were either pre-treated with neuraminidase, tunicamycin or pronase or treated with ammonium chloride before infection with H17-PV. Infectivity with H17- and H5-PVs, as well as cytotoxicity (by trypan blue exclusion), were assayed 24 h post treatment (Fig. 1d). Pretreatment of MDCK II cells with neuraminidase from *Clostridium perfrigens* (1-100 mU), which cleaves cell surface SA, reduced H5-PV infection by 68-86% but did not significantly affect infection by H17-PV, supporting the notion that SA are not the cell surface receptors for H17^19^,^26^. Classical IAVs primarily enter cells via endocytosis followed by endosomal fusion triggered by low pH. Treatment of MDCK II cells with the pH neutralising agent ammonium chloride (1-100 mM) abolished luciferase activity for both H5- and H17-PV, demonstrating that entry of H17, like IAVs, into target cells is pH-dependent. Similar results were obtained in RIE1495 cells^32^. Entry of H17-PV was more susceptible to pre-treatment of MDCK II cells with proteases or tunicamycin (an inhibitor of *N*-glycosylation) than was entry of H5-PV (being reduced by up to 72 and 78%, compared to 45 and 20%, respectively), suggesting that the H17-cellular receptor(s) may be a glycosylated protein, in line with previous proposals^19^.

**Figure 1:**
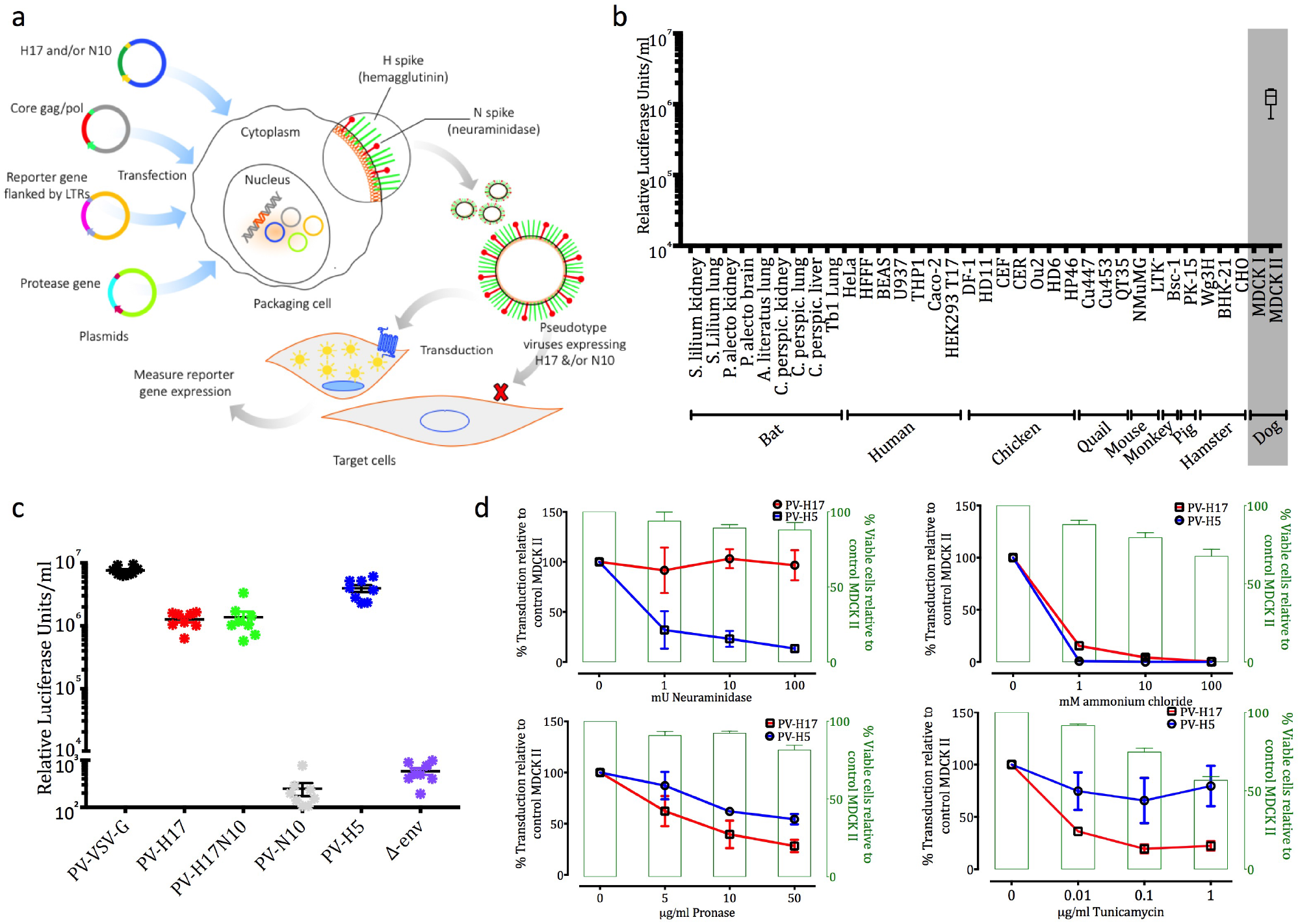
**a.** Schematic representation of pseudotype virus production. Expression plasmids for the HIV-1 gag-pol gene, the bat H17 alone or with N10, the luciferase reporter gene with HIV-1 long tandem repeats (LTRs) and the protease gene (HAT or TMPRSS2) are generated and co-transfected into producer HEK293T/17 cells. Cells transcribe and translate the HIV-1 core genes, BatIV glycoproteins are packaged on the cell surface, and viruses bud off in an HIV-1 dependent manner. Production of N10-PV does not require co-transfection with a protease gene. Supernatants are harvested at 48 h post-transfection and the produced pseudotype viruses (PVs) are filtered, titrated and infected into target cells. **b.** Infectivity titers of H17-PV in cells from different tissues/species [expressed in Relative Luciferase Units (RLU)/ml]. Mean luciferase activity was plotted as mean ± inter-assay deviation expressed as SEM from 3-6 independent experiments. **c.** Infectivity titers of H17-PV, N10 and control pseudotypes (VSV-G, H5 and Δ-env) in MDCK II cells. **d.** MDCK II cells were pre-treated with either neuraminidase (2 h), with pronase (30 mins) or tunicamycin (5 h) or treated with ammonium chloride, and then infected with H17-PV. Luciferase activities were measured after 24 h with a luminometer. The left Y-axis shows infection levels (% of control) and the right Y-axis shows % viability of cells related to control MDCK II cells. Experiments were carried out in triplicate.

MDCK I and II represent early and late passaged cells from the same parental NBL-2 cell line (CCL-34, ATCC). MDCK are valuable cell lines in studies of viruses, cell-cell junctions and epithelial differentiation but consist of heterogeneous cell populations and their phenotypes vary significantly between user laboratories^33^. In addressing the factors that permit infection of H17-PV in late passaged MDCK II cells, we considered that parental NBL-2 cells undergo passage number-dependent phenotypic changes that may be reflected at transcriptional level. The phenotypic transition of NBL-2 cells to early and late MDCK cells was investigated using the Affymetrix canine microarray 2.0 (E-GEOD-14837; passages 8 and 21 respectively). The microarray analysis identified 17 differentially regulated transcripts: 12 up-regulated and 5 down-regulated in late-compared to early-passaged cells (Fig. 2a). The current prevalent hypothesis is that single or multiple cell surface molecules are essential for the initial attachment and uptake of enveloped viruses into cells^34^, ^35^. Therefore, we surveyed the differentially regulated transcripts for encoded, surface-anchored proteins using a combined analysis of: available Gene Ontology annotations, existing literature as well as transmembrane protein domain and subcellular localisation prediction algorithms (Phobius, TMHMM and DeepLoc). The analysis identified the dog leukocyte antigen class II DR α-chain (DLA-DRA) as the only transcript, encoding a membrane protein, over-expressed in late, compared to early, passage MDCK cells (full data in Supplementary material 2). Significant over-expression of DLA-DRA (and its paralogue DLA-DRB1) was also confirmed (by qRT-PCR) in MDCK II compared to MDCK I cells (Fig. 2b).

**Figure 2:**
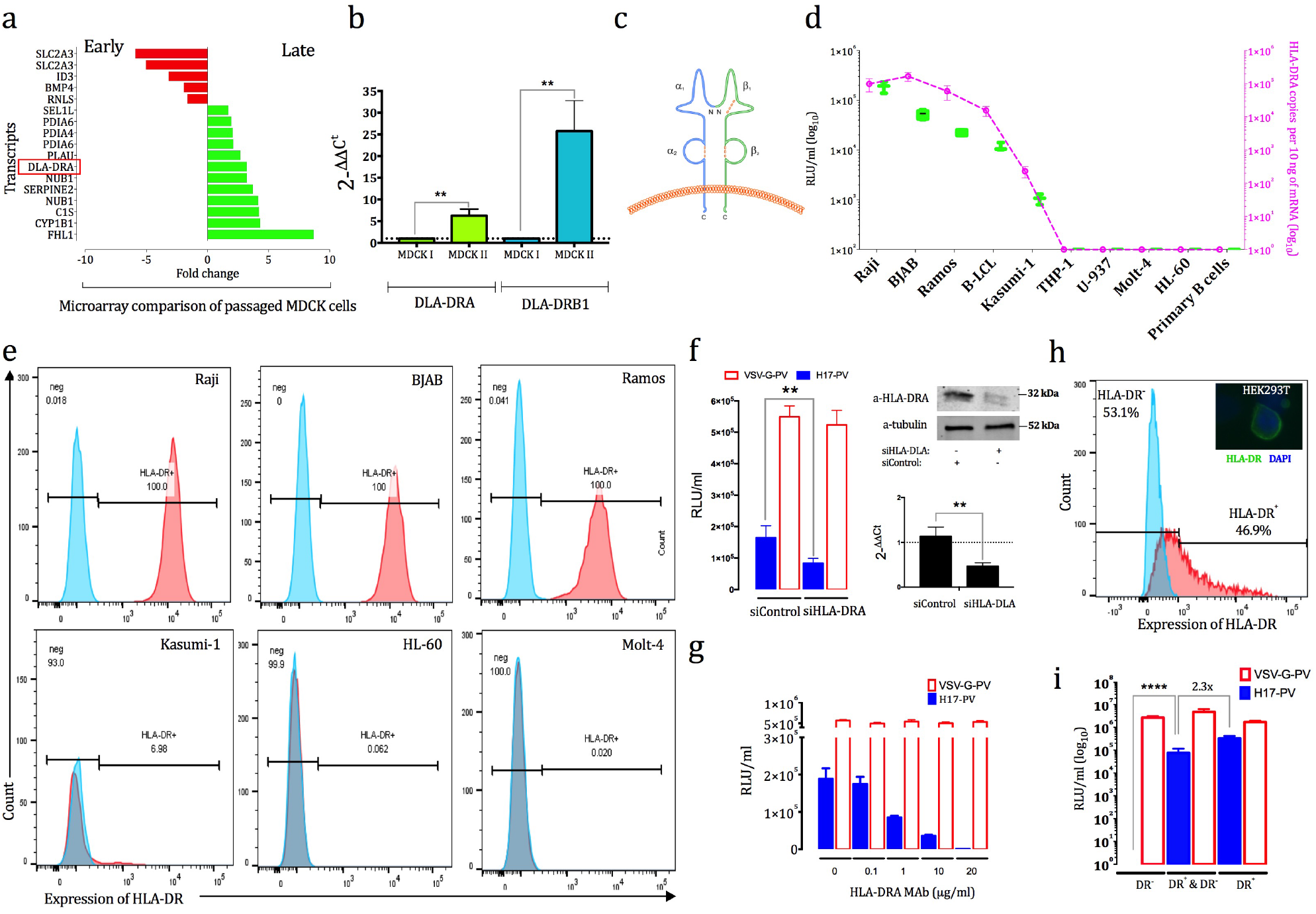
**a.** Significantly expressed genes in the microarray comparison of late versus early passaged MDCK cells (green and red columns represent upregulated transcripts in MDCK II and MDCK I cells respectively). Analysis was conducted with Partek (fold change ≥ 1.5 and FDR ≤ 0.05)). Red frame indicates the transcript upregulated in late passaged MDCK cells encoding a membrane protein. **b.** qRT-PCR results showing the expression of DLA-DRA and DLA-DRB1 mRNA in MDCK I and II cells. The data are expressed as the means ± S.E.M. from three independent experiments. One-way Anova with Tukey *posthoc* test were used to analyse the data. \*\**P*<0.005 versus control. **c.** Schematic diagram of the MHC II surface molecules. **d.** Left Y-axis (green box and whiskers): relative infection titers of H17-PV [log_10_ Relative Luciferase Units (RLU) /ml] in a panel of human cancer cell lines. Right Y-axis (broken purple line) shows log_10_ HLA-DRA mRNA copies with qRT-PCR. The data are expressed as the means ± S.E.M. from three independent experiments. **e.** FACS analysis of the expression levels of cell-surface HLA-DR molecule from three H17-PV susceptible and three unsusceptible cancer cell lines. Results are representative from two independent experiments. Blue and read peaks represent HLA-DR^-^ and HLA-DR^+^ subpopulations respectively. **f.** Left: infection titers of VSV-G- and H17-PV (RLU/ml) in Raji cells transfected with siControl or siHLA-DRA. Right: western blot (top) and qRT-PCR (bottom) showing expression of HLA-DRA protein and mRNA in transfected Raji with siRNA versus siControl. Experiments were carried out twice and the data are expressed as means ± S.E.M. One-way Anova with Dunnett *posthoc* test were used to analyse the data. \*\**P*<0.005 versus siControl. **g.** Relative infection titers of VSV-G- and H17-PV (RLU/ml) in Raji cells incubated with different concentrations of a monoclonal antibody targeting HLA-DRA. The data are expressed as the means ± S.E.M. from three independent experiments. **h.** FACS analysis of the expression levels of cell-surface HLA-DR heterodimer in transiently transfected HEK293T/17 cells for 48 h with expression vectors for DRA and DRB1 (1:1 ratio). Right hand corner microscopy picture (80x) shows immunofluorescence confirming surface expression of HLA-DR on cells (green stain: FITC-HLA-DR and blue: DAPI). The data are representative from two independent experiments. **i.** Relative infection titers of VSV-G- and H17-PV [in log_10_ Relative Luciferase Units (RLU) /ml] in FACS-sorted DR^-^ HEK293T/17 cells, unsorted transiently transfected (DR^+^ & DR^-^) and FACS-sorted DR^+^ cells. Experiments were carried out twice and the data are expressed as the means ± S.E.M. Oneway Anova with Dunnett *posthoc* test were used to analyse the data. \*\**P*<0.005 versus DR^-^ cells.

DLA-DRA is a well-conserved orthologue of the human leukocyte antigen class II DR α-chain (HLA-DRA) (~90% amino acid identity between canine, human and *Desmodus rotundus* bat ectodomains; Supplementary material 3). In humans, MHC-II molecules occur as three highly polymorphic isotypes (HLA-DR, HLA-DP and HLA-DQ) which are selectively expressed under normal conditions on the surface of antigen presenting cells (APCs), including B, dendritic and mononuclear phagocyte cells. These molecules are non-covalently associated heterodimers of two glycosylated, transmembrane polypeptide chains, the monomorphic 35-kDa α-chain and the highly polymorphic 28-kDa β-chain^36^. Both chains have an extracellular portion composed of two domains (α1 and α2, or β1 and β2) that is anchored on the cell membrane by short transmembrane and cytoplasmic domains (Fig. 2c). In the classical scenario, the protease-derived foreign antigen peptides bind to MHC class II proteins in the cleft formed by the α1 and β1 domains, and the complex is transported to the cell surface^36^, ^37^. When antigenic peptides are not available, endogenous peptides such as the class II associated invariant peptide (CLIP) substitute them and restore MHC class II dimer stability^37^. The complex of HLA-DR and endocytosed peptides (usually 9-30 amino acids in length), constitutes a ligand for the T-cell receptor (TCR) and plays a key role in the presentation of foreign antigens to CD4^+^ T helper cells and immune surveillance^38^, ^39^.

Since the ultimate goal of our studies is to obtain insight on the zoonotic potential of H17N10, we focused on the possible influence of HLA-DR on cellular susceptibility to H17. Hence, taking advantage of the high expression of HLA-DR on certain human hematopoietic carcinomas^40^, we further explored H17-PV tropism using a panel of human leukaemia and lymphoma cell lines (Fig. 2d). We found that the Burkitt’s lymphoma-derived Raji, Ramos and BJAB B-lymphocytes and the B lymphoblastoid cells (B-LCL) show decreasing susceptibility, in that order, to H17-PV. Kasumi-1 leukaemic cells showed marginal susceptibility in terms of luciferase activity, while Molt-4 and HL-60 leukaemic cells, Jurkat T-cells, pro-monocytic THP-1 and U-937 cells, and primary B cells showed no susceptibility to the pseudotypes (Fig. 2d, left Y axis). We hypothesised that the different susceptibility of the various cell types by H17-PV were due to disparate expression of HLA-DR, confirmed by qRT-PCR analysis for a non-polymorphic region of the α chain of HLA-DR on the same samples (Fig. 2d, right Y-axis). The presence of the HLA-DR heterodimer was also confirmed by flow cytometry with a FITC-conjugated monoclonal antibody (clone Tü36), which specifically binds to a monomorphic epitope on the HLA-DR α/β complex and not the isolated α or β chains (Fig. 2e)^41^. Both approaches indicate association between mRNA levels, the surface expression of HLA-DR heterodimer and susceptibility to H17-PV. Raji, BJAB and Ramos B cells were found to consist of 100% HLA-DR^+^ cells; Kasumi-1 demonstrated a 7% subpopulation of HLA-DR^+^ cells and MOLT-4 and HL-60 cells were 100% HLA-DR^-^. These results were relatively constant and did not change with factors such as cell density or passage number.

To confirm the influence of HLA-DR on H17-PV entry into B cells, HLA-DR was independently suppressed by siRNA-mediated inhibition or by antibody blocking. Raji B cells were transfected twice over 48 h with an siRNA mixture specific for HLA-DRA (siHLA-DRA) or a control siRNA (siControl) and then challenged with the H17-PV for another 48 h. The efficiency of HLA-DRA knock-down was confirmed by qRT-PCR and western Blot. The mRNA and protein expression of HLA-DRA was reduced by ≥50% in Raji cells transfected with siHLA-DRA compared to those transfected with siControl. Knocking down HLA-DRA in Raji cells correspondingly reduced the infection of H17-PV by 50% (Fig. 2f).

To determine if blocking attachment of virus to the HLA-DR ectodomain can prevent its entry, Raji cells were incubated with increasing concentrations of a monoclonal antibody (mAb Clone 302CT2.3.2) targeting a monomorphic, extracellular region of the HLA-DRA antigen (HLA-DRA epitope: amino acids 48-75). The presence of the antibody significantly reduced, in a dose-dependent manner, infection with H17-PV but not VSV-G-PV (Fig. 2g).

We next sought to ascertain whether ectopic expression of HLA-DR was sufficient to increase the susceptibility of non-APC, HEK293T/17, cells to the H17-PVs. HEK293T/17 cells were transiently transfected with the DRA expression vector, alone or in combination with DRB1. Surface expression of the α/β heterodimer was validated using immunofluorescence and flow cytometry. Expression of either DRA or DRB1 alone resulted in marginal or no increase in surface staining of HLA-DR or H17-PV infection (data not shown). In contrast, 1:1 coexpression of both DRA and DRB1 formed a functional α/β heterodimer on the cell surface in approximately 47% of the cell population (Fig. 2h and Supplementary material 4). This suggests that both α and β chains are necessary for cell-surface expression, consistent with previous studies^42^, ^43^. Transient over expression of HLA-DR in HEK293T/17 cells resulted in significant infection by H17-PV. Infection was higher by more than two orders of magnitude (Fig. 2i) in Fluorescence-activated cell sorting (FACS)-sorted cells enriched for HLA-DR. Further, expression of the human HLA-DR α and β chains in the bat *Pteropus alecto* kidney PakiTO3 cells allowed infection with H17-PV (Supplementary material 5).

Taken together, HLA-DR is shown to function as a *bona fide* entry mediator for H17 but may function with unknown factors that facilitate virus internalisation. Interaction between HLA-DR and H17 may trigger viral entry through canonical receptor-mediated endocytosis, but could also trigger entry through an activation of cell signalling pathways that the virus subverts to its advantage. Our finding therefore raises questions on the utility and possible evolutionary advantage(s) that an APC-associated receptor would confer to H17N10 infectivity and broader fitness. Some viruses exploit cells of the immune system, such as macrophages, B and dendritic cells, either as viral reservoirs or as “Trojan horses” to penetrate the epithelial barriers^44^, ^45^. The measles virus for example, exploits macrophages or dendritic cells, which traffic the virus to bronchus-associated lymphoid tissue or regional lymph nodes, resulting in local amplification and subsequent systemic dissemination by viremia^46^. A similar strategy employed by H17N10 could provide an explanation on why viral RNA was detected in different organs and tissues in carrier *S. lilium* bats (i.e. lung, kidney, liver, intestine) and why the virus fails to grow *in vitro* in cell lines developed from the same tissues^1^,^26^. The prototypic gammaherpesvirus, Epstein-Barr virus (EBV), employs resting B cells as transfer vehicles for infection of epithelial cells^47^, and also uses the HLA-DR (β1 domain) as receptor in order both to enter B cells as well as to impair antigen presentation (by sterically blocking the engagement of HLA-DR1 and the TCR Vα domain)^48^, ^49^, ^50^. It is possible that through efficient binding to HLA-DR, H17N10 may have developed a means of simultaneously accessing lymphoid cells and blocking T-cell responses. Such an immune evasion mechanism could explain, at least partially, its survival and asymptomatic status in carrier bats. With limited functional information available on the bat MHC-II, the biological role of the host HLA-DR orthologue in the pathogenesis and transmission mechanisms of the H17N10 virus remains obscure.

In this study we did not establish the stoichiometry of the HLA-DR: H17 engagement, or clarify how the virus moves to sub-membranous regions and might hijack the receptor-mediated signalling pathway to promote its internalization. It is likely that the determinants of viral entry *in vivo* are more complicated. We cannot rule out the use by H17N10 (as by other bat-borne viruses, *e.g*. SARS CoV^51^, ^52^) of more than one molecular species as (co-) receptors. Nevertheless, the implication of this study is that H17N10 has the capacity to enter human HLA-DR^+^ cells and our work provides substantial evidence that the H17N10 virus has zoonotic potential. The current finding not only sheds light on the understanding of BatIV host range, but also provides additional information on the evolution of influenza A viruses.

## Materials and methods

### Cell lines, cell culture and treatment

Cell lines (complete description in Supplementary material 1) were kindly provided as follows: the HEK293T/17 cells were provided by Dr Edward Wright (University of Westminster, UK); Kasumi-1, HL-60, Molt-4, Jurkat cells from Professor Paul Farrell (Imperial College London, UK); *Pteropus alecto* cell lines from Professor Linfa Wang (NUS Duke, Singapore); *Sturnira lilium, Artibeus planirostris*, and *Carolia perspicillata* cell lines have been generated in the labs of Dr Carles Martínez-Romero/Professor Adolfo Garcia-Sastre (Icahn School of Medicine, New York) from bat tissue samples originally collected by Dr Eugenia Corrales-Aguilar (University of Costa Rica); B-LCL were created by Dr Konstantinos Paschos by infecting with recombinant EBV B cells from isolated peripheral blood monocytes (PBMCs) of a healthy donor of the prototypical B95-8 background (Imperial College London); Raji, Ramos and BJAB from Dr Rob White (Imperial College London); U-937 and THP-1, BEAS-2B, Caco-2 from Dr Marcus Dorner, Dr Michael Edwards and Professor Robin Shattock respectively (Imperial College London). The rest of cell lines were either from the collection of Dr Michael Skinner or from ATCC. Primary B cells were a kind gift by Dr Rob White (Imperial College London). B cells were isolated from peripheral blood leukocyte (PBL) samples obtained from anonymous buffy coat donors (UK Blood Transfusion Service) by centrifugation over Ficoll. CD19 microbeads were used for magnetic separation of purified B cells using an autoMACS separator (Miltenyi Biotec). All cell lines in this study were cultured according to standard mammalian tissue culture protocols (ATCC; www.atcc.org). Bat cell lines were propagated in Dulbecco’s modified eagle medium (DMEM) (Life Technologies) supplemented with heat-inactivated 15% fetal bovine serum (Life Technologies), penicillin (100 U/ml) and streptomycin (100 μg/ml; Invitrogen). All cells were maintained in a humidified incubator at 37°C and 5% CO_2_ and were found free of mycoplasma contamination on repeated testing with the MycoFluor Mycoplasma Detection Kit (Life Technologies, UK). MDCK II cells were treated as previously^19^ with the following modifications. MDCK II cells were either treated with the endosomal acidification reagent ammonium chloride (1, 10 or 100 mM), or pre-treated with neuraminidase from *Clostridium perfrigens* for 2 h (1, 10 or 100 mM) or pronase (a mixture of endo- and exoproteases from *Streptomyces griseus* at a final concentration of 5, 10 or 50 μg/ml; Calbiochem, UK) for 30 min, an N-glycosylation inhibitor (tunicamycin from *Streptomyces* sp. at a final concentration of 0.01, 0.1 or 1 μg/ml; Sigma-Aldrich, UK) for 5 h. Pre-treated cells were washed with phosphate buffer saline (PBS) 3 times, and then incubated/infected as before with PVs for another 24 h. Cell viabilities were assessed by a trypan blue exclusion test.

### Lentiviral pseudotype virus production and susceptibility assays

Pseudotypes expressing H17 and N10 genes were produced as described previously^32^, ^53^. Briefly, the lentiviral packaging plasmid p8.91^54^, the pCSFLW firefly luciferase lentiviral vector^55^ or the GFP expressing vector pCSGW, the expression plasmids for H17 and/or N10 [vector pI.18^56^ and the protease encoding plasmid pCAGGS-HAT (a kind gift by Eva Böttcher-Friebertshäuser, Philipps University of Marburg, Germany) were co-transfected using polyethylenimine transfection reagent (Sigma Aldrich, UK) into HEK293T/17 cells, plated on 6-well Nunclon© plates (Thermo Fisher Scientific, UK). Supernatants were collected 48-72 h post transfection and filtered through a 0.45 μm filter (Millipore, UK). To remove viral titer bias between different PV stocks, pseudotypes were concentrated and (re-) titrated by serial dilution. Concentration was carried out by ultra-centrifugation for 2 h at 25,000 rpm, 4°C in the SW32 rotor of a L2-65B Beckman ultra-centrifuge.

Two-fold serial dilutions of PV-containing supernatant were performed as previously described^32^ using white 96-well Nunclon^©^ plates (Thermo Fisher Scientific, UK). Subsequently, approximately 1×10^4^ (for adherent) and 3 x10^4^ cells (for suspension) cells were added in 50 μl of medium per well. Plates were incubated for 48 h, after which 50 μl of Bright-Glo™ substrate (Promega, UK) was added. Luciferase readings were conducted with a luminometer (FLUOstar OPTIMA, BMG Labtech) after a 5-minute incubation period and luciferase reading recorded in relative luminescence units (RLU). Data were normalized using Δ-env and cell-only measurements and expressed as RLU/ml.

### Plasmids and transfections

Mammalian expression plasmids (pcDNA3.1^+^/C^-^(K)DYK) for HLA-DRA (NM_019111) and HLA-DRB1 (NM_001243965) were purchased from GenScript (Piscataway, NJ; USA). HEK293T/17 cells at sub-confluence in 6-well plates or 100 mm dishes were transfected with HLA-DRA plasmid or HLA-DRB1 plasmid or a 1:1 combination of both using the Lipofectamine 3000 transfection reagent (Thermo Fisher Scientific, UK) according to manufacturer instructions. 48 h after transfection, cells were used either for immunofluorescence analysis, or for FACS analysis (cell-surface staining) or for infection with PV. Under the experimental conditions, the transfection efficiency in either plate/dish, as assessed by the GFP expression of a co-transfected GFP-expressing control plasmid, was >70% under microscopic observation. For HLA-DR stably over expressing cells, PakiTO3 cells were transfected with both HLA-DR plasmids and then selected with neomycin (500 μg/ml). Single clones were analysed for expression of the over expressed proteins.

### RNA isolation for microarray analysis

Total RNA was isolated from biological triplicates of early and late passage MDCK from T25 flasks that had been seeded with 8 x 10^5^ cells and allowed to become confluent and polarize over 4 days in culture cells using a Ribopure kit (Ambion, Austin, TX). Acquired RNA was precipitated with EtOH and subsequently purified employing columns, procedures and reagents from an RNEasy kit (Qiagen, Germantown, MD) and resuspended in RNAse-free H_2_O. Complementary DNA and RNA synthesis were performed according to Affymetrix Expression

Analysis protocols (see www.affymetrix.com). Briefly, double-stranded cDNA was synthesized from 5 μg of total RNA using the Superscript double-stranded cDNA synthesis kit (Invitrogen). Following phenol/chloroform extraction and ethanol precipitation, a biotin-labeled in-vitro transcription reaction was carried out using the cDNA template (Enzo Life Sciences, Farmingdale, NY). Fifteen micrograms of cRNA was fragmented for hybridization to Affymetrix Canine Genome 2.0 Array GeneChips (Santa Clara, CA), which contains approximately 18,000 *C. familiaris* mRNA/EST-based transcripts and over 20,000 non-redundant predicted genes. An one-way ANOVA adjusted with the Benjamini–Hochberg multiple-testing correction [false discovery rate (FDR) of P<0.05] was performed with Partek Genomics Suite (v6.6 Partek) across all samples as previously^57^. Principal component analysis confirmed that batch mixing had prevented introduction of experimental bias. Comparisons were conducted between early and late passaged cells. The analysis cut off criteria were fold change ≥ ±1.5 and *P*-value ≤ 0.05. Microarray data was uploaded per MIAME standards and deposited at the GEO repository and is available under series record number GSE14837.

### HLA-DRA knockdown and blocking using siRNA and monoclonal antibodies

A Sigma-Aldrich MISSION esiRNA endonuclease-derived mixture of siRNAs (EHU226621) was used to knock down HLA-DRA expression in Raji cells. Lipofectamine RNAiMAX transfection Reagent (Thermo Fisher Scientific, UK) was used to transfect exponentially grown Raji cells with 50 nM of siHLA-DRA or siRNA universal negative control (Sigma-Aldrich, UK; SIC001) according to the manufacturer’s instructions. The transfection was repeated the following day and cells were collected after 48 h and either seeded at 3×10^4^ cells per well in a 96 well-plate for infection with PV or processed in order to validate siRNA activity. Total RNA and protein were collected and assessed by quantitative RT-PCR and western blot, respectively.

In order to evaluate the interaction of HLA-DR with H17 we used the HLA DRA mAb (302CT2.3.2), which is generated from mice immunized with a KLH conjugated synthetic peptide between 48-75 amino acids from human HLA-DRA. After a 1 h pre-incubation with increasing concentrations of the antibody in normal growth media, Raji cells (3×10^4^) were infected for 24-48 h with H17-or VSV-G-PV.

### Western blot analysis

Washed cells were lysed on ice with lysis buffer [0.5% NP40 in PBS with 10 mM Tris-HCl, pH 7.4 supplemented with Halt Protease Inhibitor mixture EDTA-free (Thermo Fisher Scientific UK)] and protein was quantified by the BCA assay kit (Thermo Fisher Scientific, UK). 20-50 μg of protein was electrophoresed on a 4-15% sodium dodecylsulfate polyacrylamide gel, alongside a protein ladder (Precision Plus Protein Dual Colour Standards, Bio-Rad) and immunoblotted with the following antibodies using standard procedures: mouse monoclonal anti-HLA-DRA (1:1000; Clone: 302CT2, Enzo Life Sciences, UK) or rabbit monoclonal a-tubulin (1:2000; Cell signalling Technology, UK) antibodies. The membranes were then washed with PBS for three times and incubated with goat anti-rabbit or donkey anti-mouse secondary antibodies (LI-COR) in the dark for 1 h. Scanning was then carried out using the Odyssey Imaging system (LI-COR).

### Immunofluorescence

Transfected cells with HLA-DRA and -DRB1 expression plasmids or with the empty plasmid were seeded onto glass cover slips at 5×10^4^ cells/ml in 6 well plates overnight and were fixed with 4% paraformaldehyde in PBS for 30 minutes at room temperature (RT). Fixed cells were washed with PBS and permeabilised with 1% Triton X-100 in PBS for 10 minutes. After washing with PBS, the cover slips were incubated with a mouse HLA-DRA mAb (169-1B5.2; Bio-Techne Ltd) targeting a monomorphic general framework determinant of HLA-DR Class II antigen, diluted in 5% BSA/PBS for 1 hr at RT. The cover slips were then washed 3X with 0. 02% Tween 20 and 1% BSA in PBS, followed by incubation with Alexafluor 488 conjugated anti-mouse (Thermo Fisher Scientific, UK) for 30 minutes at RT. After washing 3X with 0.02% Tween 20 and 1% BSA in PBS, the cover slips were mounted using Prolong Gold containing DAPI (Invitrogen). Images were acquired on EVOS fluorescent microscope (EVOS FL imaging system; Life Technology, USA). Experiments were carried out twice.

### Flow cytometry

For surface staining, human cancer cells and HEK293T/17 cells transfected with an empty vector or expression plasmids of HLA-DR (α and β) chains were maintained in the dark at 4°C throughout. Cells were collected, washed twice in ice-cold FACS buffer (2%FCS, 0.02% NaN3 in PBS) and stained with a FITC-conjugated anti-human HLA-DR mAb (Clone Tü36; BD Biosciences). This antibody specifically binds to a monomorphic epitope of the HLA-DR αβ complex and not the isolated α or β chains or the HLA-DP and -DQ isotypes^41^. The cells were analyzed with BD LSR Fortressa to determine expression of HLA-DR in combination with the matched isotype control and sorted into HLA-DR^-^ and HLA-DR^+^ subpopulations with a FACS Aria cell sorter (BDIS, San Jose, CA, USA). The Hoechst 33342 stain was used for cell viability discrimination and the data files were analyzed using the FlowJo software (Tree Star, Inc., Sac Carlos, CA, USA). Data are representative of two independent experiments.

### Relative and absolute mRNA quantification

qRT-PCR was performed on RNA samples using a two-step procedure. RNA was first reverse-transcribed into cDNA using the QuantiTect Reverse Transcription Kit (Qiagen) according to manufacturer’s instructions. qRT-PCR was then conducted on the cDNA in a 384-well plate with a ABI-7900HT Fast qRT-PCR system (Applied Biosystems). Mesa Green qRT-PCR MasterMix (Eurogentec) was added to the cDNA (5 μl for every 2 μl of cDNA). The following primers were used: for canine DLA-DRA (Forward: 5’-GCTGTGGACAAAGCTAACCTTG-3’, Reverse: 5’-TCTGGAGGTACATTGGTGTTCG-3’), for canine DLA-DRB1 (Forward: 5’- AGCACCAAGTTTGACAAGC-3’, Reverse: 5-AAGAGCAGACCCAGGACAAAG-3’). The following amplification conditions were used: 95°C for 5 minutes; 40 cycles of 95°C for 15 seconds, 57°C for 20 seconds, and 72°C for 20 seconds; 95°C for 15 seconds; 60°C for 15 seconds; and 95°C for 15 seconds. The output Ct values and dissociation curves were analysed using SDS v2.3 and RQ Manager v1.2 (Applied Biosystems). Gene expression data were normalized against the housekeeping gene GAPDH, and compared with the mock controls using the comparative Ct method (also referred to as the 2^-ΔΔCT^ method^58^). Absolute copy numbers of HLA-DRA in human cell lines were calculated using a standard curve of known concentrations of the corresponding HLA-DRA cDNA expression plasmid. HLA-DRA (Forward: 5’- TCAAGGGATTGCGCAAAAGC-3’ and reverse 5’- ACACCATCACCTCCATGTGC-3’. All experiments were carried out in triplicate.

### Prediction of transmembrane protein domains and subcellular topology and phylogenetic analysis

For each of the differentially regulated transcripts identified by microarrays, we used Phobius (http://phobius.sbc.su.se)^59^ and TMHMM v2.0 (http://www.cbs.dtu.dk/services/TMHMM/)^60^ to predict the existence of transmembrane protein domains. Similarly, we used Deeploc-1.0 (http://www.cbs.dtu.dk/services/DeepLoc/)^61^ to determine sub-cellular localisation of the encoded proteins. These predictions were compared with gene annotations and literature references to confirm their reliability. The amino acid sequences of canine DLA-DRA (NP_001011723.1), human HLA-DRA (NP_061984.2), and their bat orthologues [(*Pteropus alecto* (XP_006907484.1) and *Desmodus rotundus* (XP_024413747.1)] were subjected to multiple alignment using CLC workbench 7 (CLC Bio, Qiagen, Aarhus, Denmark).

### Ethics statement

The buffy coat residues for the isolation of CD19^+^ primary B cells were purchased from the UK Blood Transfusion Service from anonymous volunteers blood donors. Therefore, no ethical approval is required.

### Statistical Analyses

Graphical representation and statistical analyses were performed using Prism 8 software (GraphPad). Unless otherwise stated, results are shown as means ± SEM from three independent experiments. Differences were tested for statistical significance using one-way ANOVA with a Dunnett’s or a Tukey *posthoc* test. All statistical analyses were two-sided, and p <0.05 was considered statistically significant.

## Acknowledgements

Our thanks go to Jim Kaufman, Yanping Guo, Rob White, Amr Bayoumy, Ibrahim Elbusifi, Daragh Quinn and Alfred Ho for their technical assistance. This research was undertaken with the financial support of the Biotechnology and Biological Sciences Research Council (BBSRC) (http://www.bbsrc.ac.uk) via Strategic LoLa grant BB/K002465/1 “Developing Rapid Responses to Emerging Virus Infections of Poultry (DRREVIP)” and the Octoberwoman Foundation.

## Supplementary material

**Supplementary Material 1:**
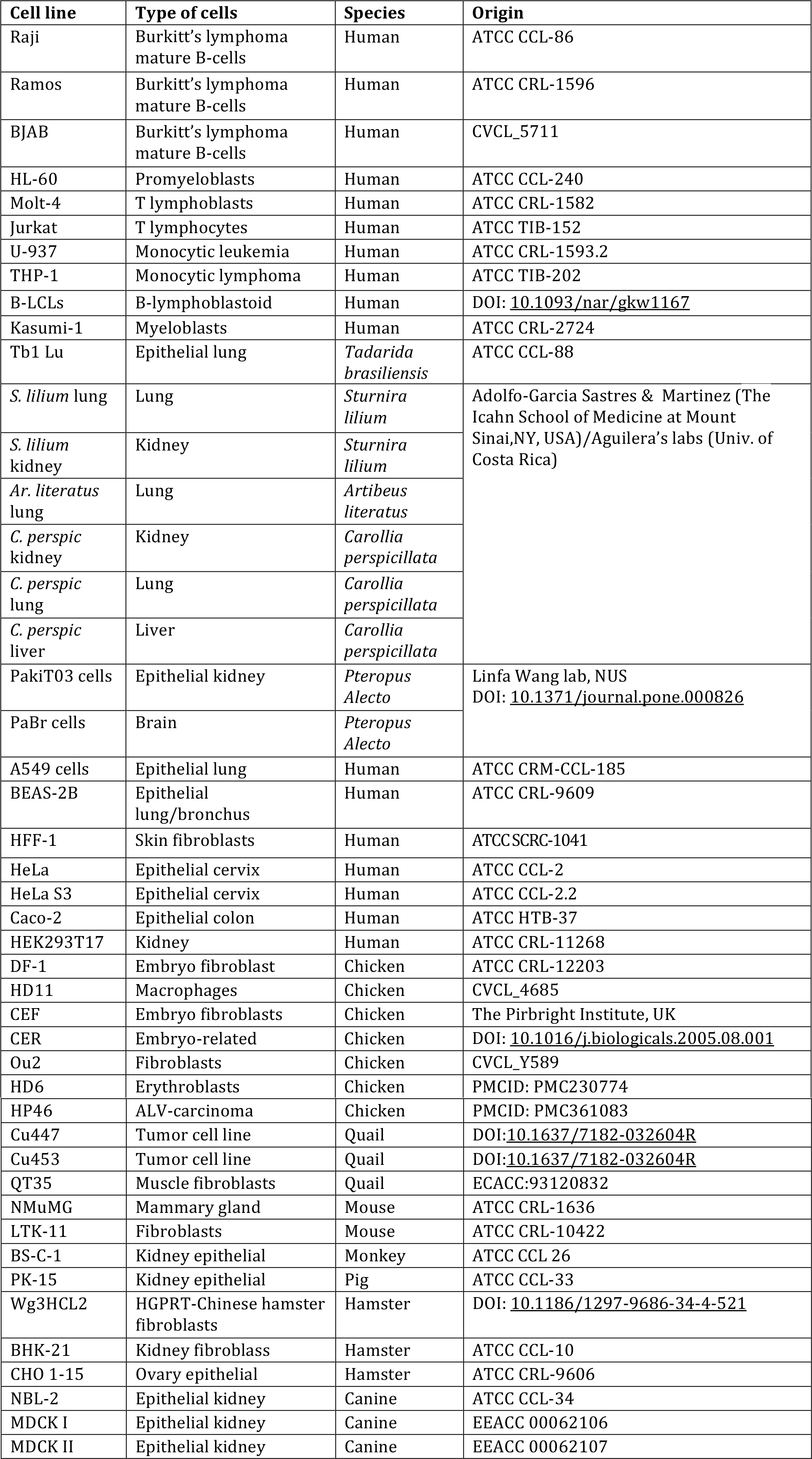
List of cell lines included in the study.

**Supplementary Material 2:**
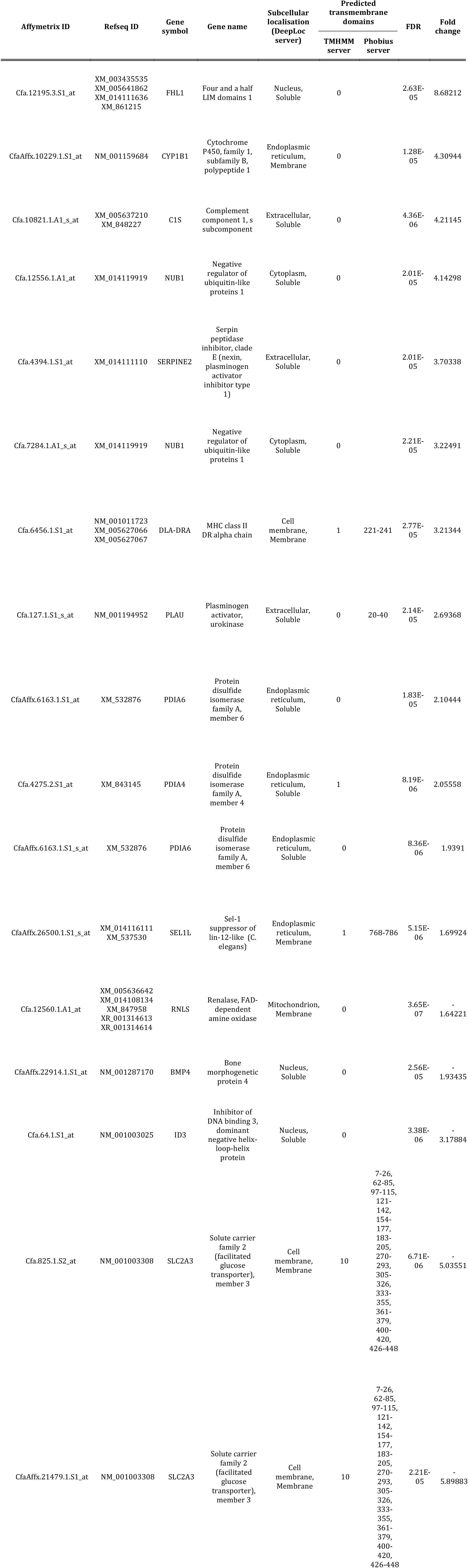
List of the differentially regulated genes determined by microarray comparison between early and late passaged MDCK cells, as summarised in Fig. 2a. The encoded proteins of the differentially regulated transcripts were surveyed for their subcellular localisation (with the DeepLoc server) and presence of transmembrane domains (using the PHOBIUS and TMHMM algorithms).

**Supplementary Material 3:**
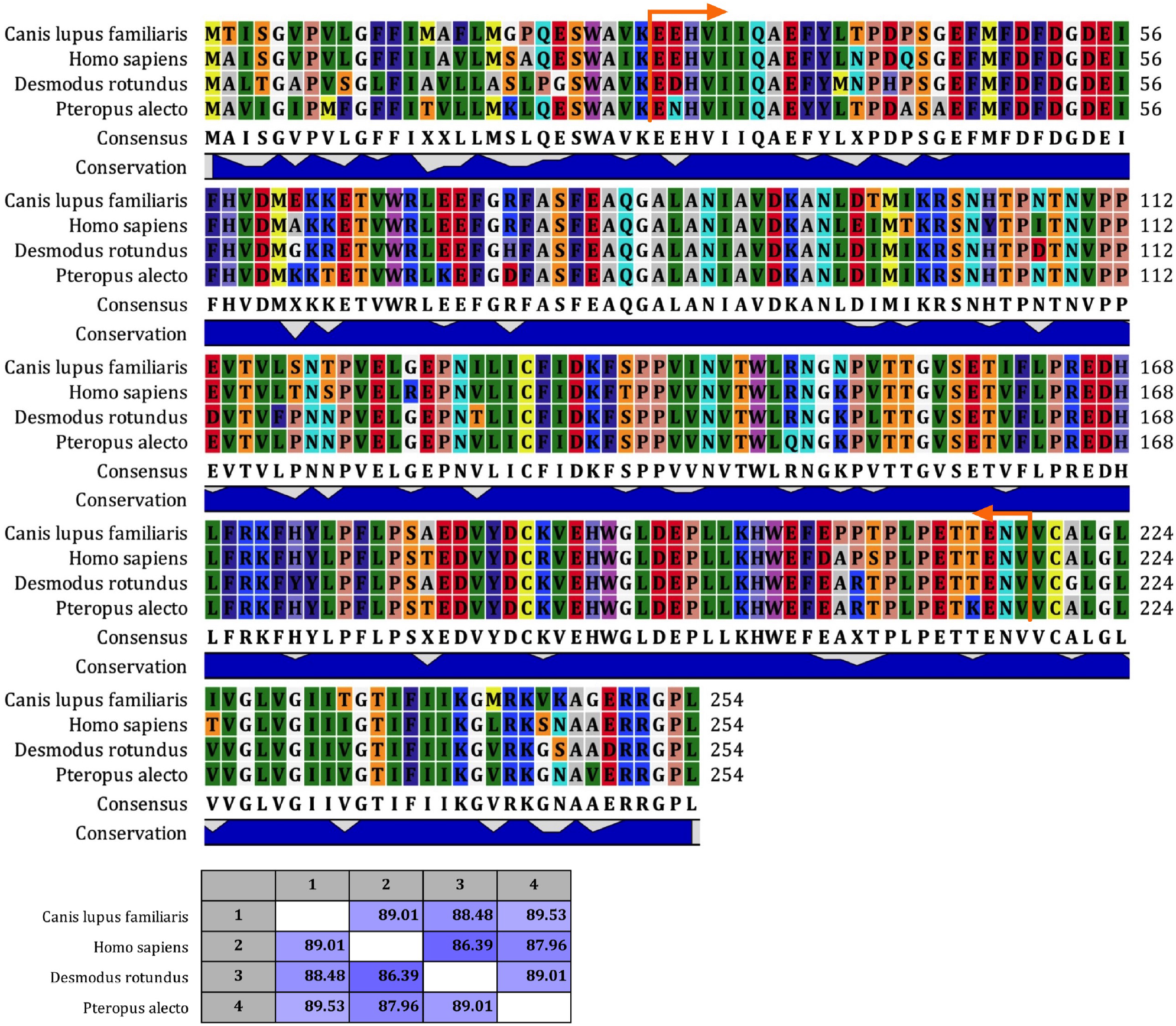
The ectodomain of HLA-DRA is well conserved in bats, humans and canine. Multiple amino acid sequence alignment of canine DLA-DRA (NP_001011723.1), human HLA-DRA (NP_061984.2), and their bat orthologues [(Yinpterochiroptera *Pteropus alecto* (XP_006907484.1) and Yangochiroptera *Desmodus rotundus* (XP_024413747.1)]. *Desmodus rotundus* (common vampire bat) was chosen as the closest species to *Sturnira lilium* with a decoded genome. The alignment was performed by importing the corresponding amino acid sequences into CLC Workbench (CLC Bio, Qiagen, Aarhus, Denmark). Orange arrows indicate the ectodomain of the protein. Matrix below shows overall, pairwise amino acid similarity of HLA-DRA between the four species. The percent amino acid similarity values were calculated with the CLC workbench program.

**Supplementary Material 4:**
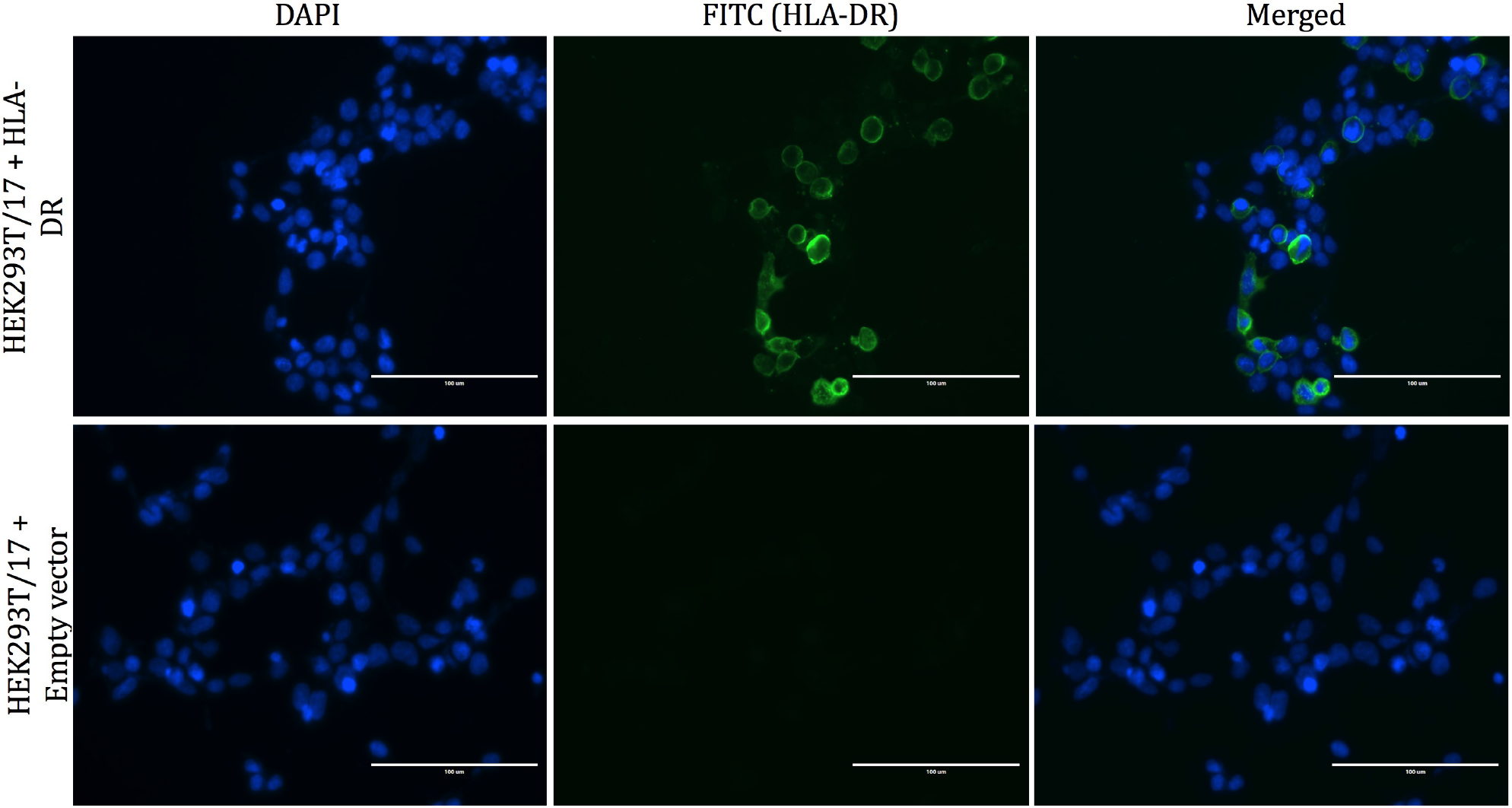
Transfection of HEK293T/17 with HLA-DR α and β chains results in surface expression of the heterodimer. HEK293T/17 cells were transfected with the empty vector or with HLA-DRA and DRB1 expression plasmids. The cells were fixed in paraformaldehyde and were permeabilised with Triton X-1OO to show intracellular distribution and immuno-stained with mAb HLA-DRA. Nuclei were stained blue with DAPI stain (left panel), HLA-DR heterodimers were stained green (middle panel), and the right panel shows a merged image. Fluorescent microscopy analysis was performed with the EVOS FL fluorescent imaging system. Original magnification, × 20.

**Supplementary Material 5:**
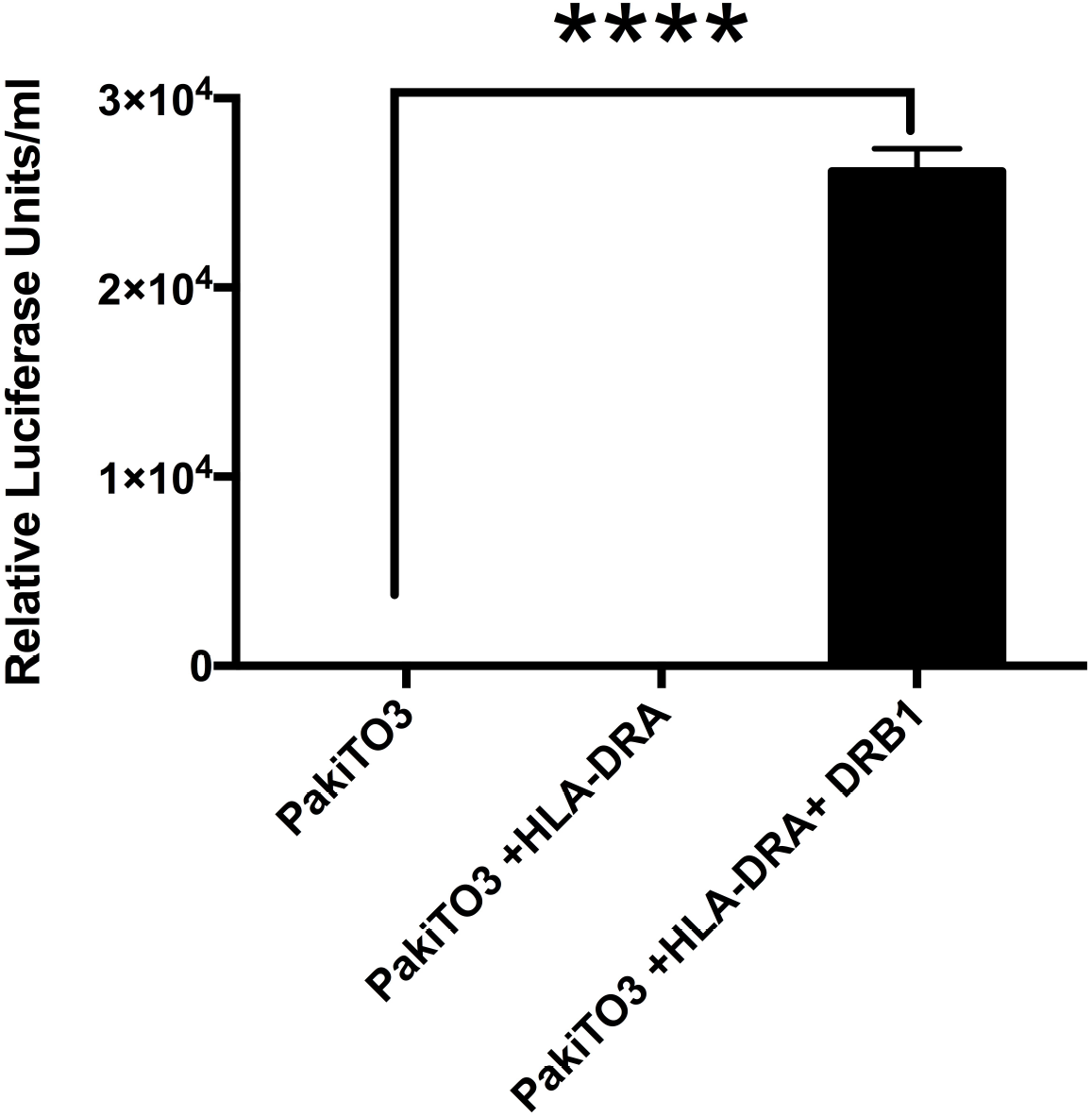
Transfection of bat PakiTO3 cells with HLA-DR α and β chains confers susceptibility to H17-PV. Infectivity titers of H17-PV (RLU/ml) in PakiTO3 cells transfected for 48 h with equimolar amounts of either empty vector (pcDNA3.1) or the expression plasmid for HLA-DRA or the expression plasmids encoding both chains of HLA-DR. Cells were infected with PV for an extra 72 h in the presence of neomycin. Data represent mean values ± SEM of three independent experiments. One-way Anova with Dunnett *posthoc* test were used to analyse the data. \*\*\*\**P*<0.00005 versus cells transfected with the empty vector.

## Abbreviations

SARS: severe acute respiratory syndrome
MERS: Middle-East respiratory syndrome
CoV: coronavirus
MDCK: Madin-Darby canine kidney cells
HIV: human immunodeficiency virus
CD4: cluster of differentiation 4
MHC: Major histocompatibility complex
SARS: severe acute respiratory syndrome
ATCC: American type culture collection
gag: group-specific antigen
pol: polymerase
SEM: standard error of the mean.
SARS: severe acute respiratory syndrome

